# Mitochondrial DNA and the 641kb nuclear-mitochondrial DNA in *Arabidopsis* can be separated by their CpG methylation levels

**DOI:** 10.1101/2024.05.14.594123

**Authors:** Yuyang Zhong, Miki Okuno, Nobuhiro Tsutsumi, Shin-ichi Arimura

## Abstract

Methylation on the cytosine in plant mitochondrial DNA has been a controversial issue. Results supporting mitochondrial DNA methylation may have been subject to contamination due to the presence of the nuclear sequence originating from the mitochondrial genome called NUMT (nuclear mitochondrial DNA insertions). Especially in *Arabidopsis thaliana* Columbia 0, the largest NUMT located on chromosome 2 is nearly twice the size of the entire mitochondrial genome and almost identical. In the presence of such high similarity, it is challenging to eliminate interference in the determination of mitochondrial DNA methylation levels. Here we applied MBD (methyl-CpG binding domain) protein-based affinity assay to separate total DNA, and applied next generation sequencing to the pre- and post-separation DNA samples, and checked their SNP sites. The results revealed successful separation of methylated and non-methylated DNA within the total DNA, with separation achieved between NUMT DNA and mitochondrial DNA. The result suggests that our method can achieve separation based on the differential methylation levels of the whole lengths of NUMT and mitochondrial DNAs, and that mitochondrial DNA barely exhibits CpG methylation, at least in the Columbia 0.

## Introduction

As one of the epigenetic modifications, DNA methylation plays a crucial role in the physiological processes of organisms (Moore *et al*., 2013, Zhang *et al*., 2018). Mitochondrial DNA (mitoDNA) methylation of mammals, fungi, and plants has been explored so far (Kang *et al*., 2017, Muniandy *et al*., 2020, Shock *et al*., 2011). The mitoDNA methylation level of *Arabidopsis thaliana* (*A*.*thaliana*), ecotype Columbia 0 (Col-0) has been observed to exist and decrease in *demethylation regulator 1* (*demr 1*) mutant through comparison of changes in whole-genome bisulfite sequencing (WGBS) (Wang *et al*., 2022). It is worth noting that the presence of NUMT (nuclear mitochondrial insertion) was not taken into account in this paper. Located on the nuclear genome, the NUMT, in some cases, shares a high similarity with the mitochondrial genome (Wei *et al*., 2022, Noutsos *et al*., 2005), which may be methylated to some extent and thus interfere with the determination of mitochondrial DNA methylation. Especially in the case of *A. thaliana* Col-0, the largest NUMT of which is located on the chromosome 2. Not only is this NUMT larger than the entire Col-0 mitochondrial genome (641kb vs 367kb), but it also shares 99.933% sequence identity with the mitochondrial genome (Fields *et al*., 2022a). The commonly used methods for detecting DNA methylation are bisulfate conversion-based methods such as whole genome bisulfate sequencing (Gu *et al*., 2011), affinity enrichment-based methods such as methylated DNA immunoprecipitation (MeDIP or mDIP) (Zhao *et al*., 2014), and restriction enzyme-based methods such as <u>c</u>omprehensive <u>h</u>igh-throughput <u>a</u>rrays for relative <u>m</u>ethylation (CHARM) (Irizarry *et al*., 2008). These assays often require the digestion of long sequences into shorter sequences before they can be read and detected. Using any of these methods, if NUMT presents, methylation of DNAs from NUMT is highly likely to be counted as methylation of mitoDNA. Even for bisulfate sequencing serves as the gold standard for cytosine methylation measurement, there’s still the issue of incomplete conversion of unmethylated cytosine to uracil (Matsuda *et al*., 2018). Moreover, some research findings challenge the perspective on human and mouse mitoDNA methylation (Bicci *et al*., 2021, Mechta *et al*., 2017). Although NUMT poses a significant interference to the determination of mitoDNA methylation, it is fortunate that high-quality NUMT sequence information have already been published, indicating the biggest NUMT on chromosome 2 in Col-0 is highly methylated at 5mC (Fields et al., 2022a). In this research, we aim to investigate whether the mitochondrial DNA of Col-0 is CpG methylated by circumventing NUMT-induced interference in the determination of mitoDNA methylation.

## Results

### The CpG methylation status of NUMT and mitochondrial sequences by using SNPs between them

Plant DNA cytosine methylation occurs on CG (55 %), CHG (23 %), CHH (22 %) contexts in nucleus (Lister *et al*., 2008). In this study, we focus on the CG context in mitochondrial DNA. CpG-methylation sensitive restriction enzyme (RE) *Pvu*I and CpG-methylation dependent RE McrBC (Sutherland *et al*., 1992) have been used in this case. Total DNA from leaves of Col-0 (1-month-old) was extracted and digested by these two REs first, and then amplification was applied on a *rps3* gene area shared by both mitochondrial and NUMT sequences. Sanger sequencing to see the SNP at +1904C/T in *rps3* between mitochondrial and NUMT sequences was applied to the amplification products (Fig.1a). In this RE and amplification assay, for both non-restriction enzyme treatment controls, the overlap of NUMT-type SNPs (numtSNPs) and mitochondrial-type SNPs (mitoSNPs) in the Sanger sequencing chromatogram peaks was observed. After the McrBC treatment, only mitoSNP remains, suggesting that all the NUMT *rps3* sequences seemed to be digested, and mitochondrial DNA remained to be amplified. As for *Pvu*I treatment, only the numtSNP remains, suggesting that mitochondrial DNA in the targeted *rps3* area was digested by *Pvu*I so that it can’t be amplified. This result indicates no CpG-methylation, at least in the *rps3* area in the mitochondrial genome, but the complete CpG methylation of the corresponding NUMT *rps3* area.

**Fig. 1.**
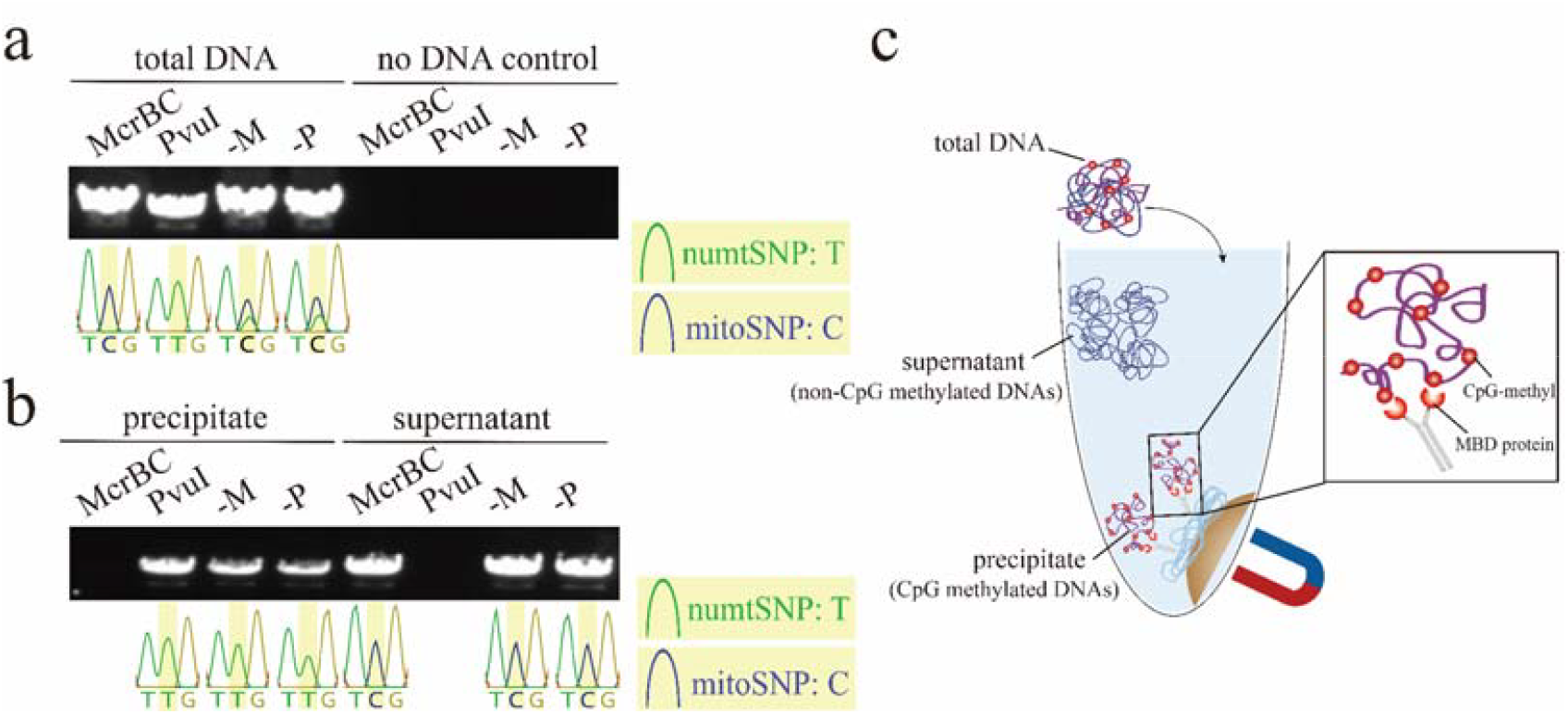
The amplification of the *rps3* in total DNA after restriction enzyme treatment (a-c) and the proportion of mito-/numtSNPs in certain genes. **a**,**b** The base in the middle (yellow shade) of the PCR-direct Sanger sequencing chromatogram peak result represents the SNP sites. McrBC, CpG-methylation dependent restriction enzyme. *Pvu*I, CpG-methylation sensitive restriction enzyme. numtSNP: the type of SNP on NUMT; mitoSNP: the type of SNP on mitochondria. -M: No McrBC control, only the buffer of McrBC was used; -P: No *Pvu*I control. **c**, the rationale of MBD protein-binding precipitation.

Next, we used an MBD (methyl-CpG binding domain) protein-based affinity precipitation assay to separate the CpG-contained DNA and others from the total DNA. The MBD protein can only bind to the CpG methylated DNA sequences (Nan *et al*., 1993), therefore, when total DNA was put into the MBD-protein involved system, the CpG methylated DNA will be captured and separated (Fig.1c). Similar RE and amplification assay was applied to the precipitate and the supernatant samples (Fig. 1b). In the precipitated sample, all the sequences were digested by McrBC, suggesting the high CpG methylation in this portion; while in the supernatant sample, all the sequences were digested by *Pvu*I, suggesting an absence of CpG methylation in this portion. Taken together, the separation of CpG-contained DNA and non-methylated DNA were achieved by this MBD protein-based assay. Furthermore, in precipitated samples, the SNPs of RE and no-RE treatment were both numtSNP, suggesting that the precipitate part contains exclusively NUMT. For the supernatant sample, the SNPs of RE and no-RE treatment were mitoSNPs, suggesting that the supernatant part contains exclusively DNAs from mitochondria. Collectively, the precipitated sample is highly CpG methylated and contains DNA from NUMT, at least in the *rps3* area, while the supernatant sample is absent from CpG methylation and contains DNA of the corresponding area from mitochondria. This result hinted the MBD protein-based affinity precipitation assay allows for the separation of mitoDNA and NUMT.

Illumina-based next-generation sequencing was applied to the total DNA, precipitated DNAs, and supernatant DNAs. The short reads from these three DNA samples were then mapped to the reference genome of NUMT and the mitochondrial genome. Their read coverages and a dot plot analysis are shown in Figure 2. In contrast to the coverage pattern of precipitated DNA mapped to NUMT reference sequence looked overall flatly at all length, the short reads from the precipitated DNA sample aligned to the mitochondrial reference genome contains two regions (in red shadow) of the three to four-times increased number of reads mapped were observed. The dot plot shows that these two regions correspond to the three (or four) times repeated sequences in the NUMT. These repetitive sequences are due to post-insertion duplications (Hazkani-Covo *et al*., 2003). This implies that almost all the reads in the precipitate portion are derived from the NUMT. Similarly, the supernatant reads were mapped roughly flatly to the mitochondrial genome, the reads from supernatant portions mapped to the NUMT reference genome contains two special areas (in green shadow) of the three to four-times increased number of reads were also observed. This suggests that the supernatant reads are almost all from the mitochondrial DNA genome sequence. The coverage patterns of total DNA showed intermediate but slightly more similar to the pattern of the supernatant, which seems reasonable because the copy number of the mitochondrial DNA is much higher than the nuclear DNA in the cells (Preuten et al., 2010). Collectively, the whole lengths of NUMT and mitochondrial DNA seem separated clearly by the MBD protein-based affinity precipitation assay.

**Fig. 2.**
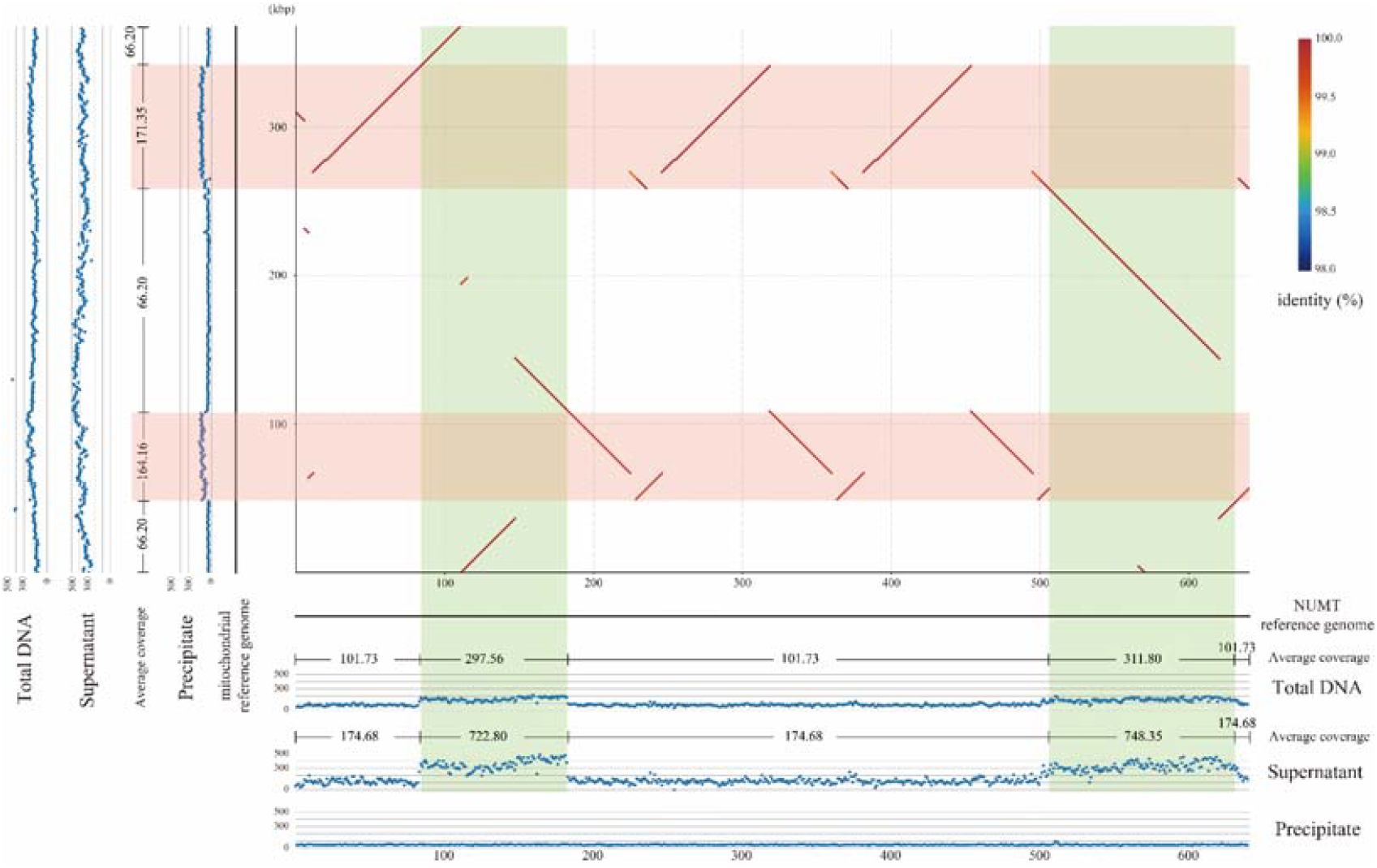
Dot-plot alignment between the mitochondrial genome and the NUMT genome, and each of the read depth distributions of the WGS data. The horizontal axis shows the NUMT region in chromosome 2 (Col-CEN) and the vertical axis shows the mitochondrial reference genome (BK010421). The lines in the dot-plot represent the aligned regions between the two sequences and colors of each line are based on sequence identity, and the color of the line (almost all in red) shows the identity (%) in the rainbow bar legend. The scatter diagrams to the left and bottom of the dot-plot show the read depth distributions of three samples mapped separately for each sequence. Multi-copy loci in the NUMT region are highlighted in red and single-copy loci are highlighted in green.

After mapping NGS reads from the three samples to the mitochondrial and NUMT genomes separately, the allele frequency (AF) of each position was calculated through dividing allelic depth (AD) by depth of coverage (DP) (Fig. 3). 246 SNP sites were found, and AF values of most SNPs are concentrated within 5% AF when mapping reads from reads of supernatant portion to the mitochondrial reference genome. 496 SNP sites were found when mapping the reads of total DNA portion to the mitochondrial reference genome, the majority of which are concentrated within the 5 ∼ 20% AF range. When mapping reads from the precipitate portion to the mitochondrial reference genome, 481 SNP sites were found and the values of AF are primarily focused within the range of 95∼100%. Compared to the other two portions, the number of SNPs after read alignment in the supernatant portion has decreased by approximately half, suggesting that the supernatant portion DNA exhibits high similarity with the mitochondrial reference genome in the full length. As for mapping to the NUMT reference sequence, 415 SNP sites were found and most of the SNPs have 0 ∼ 5% AF for the precipitate portion. 718 SNP sites were found and most of them were focused on 5 ∼ 10% AF for the total DNA portion. 636 SNP site were found and a large part of these SNPs were focused on 0 ∼ 5%, another part of these SNPs were focused on 95 ∼ 100%. Compared to the total DNA and precipitate portions, nearly half of the SNP sites was disappeared, suggesting the precipitate portion exhibits high similarity with the NUMT reference sequence. The special regions highlighted in Fig.2 are also indicated in Fig.3, where the inserted regions exhibit a greater diversity of AF values. Due to the similarity between the supernatant portion and the mitochondrial reference genome, the supernatant portion contains almost exclusively mitoDNA. As the precipitated portion resembles NUMT reference sequence, the precipitated portion contains almost exclusively NUMT DNA (numtDNA). Taken together, mitoDNAs and numtDNAs were separated at full length through the MBD protein-based affinity precipitation assay.

**Fig. 3.**
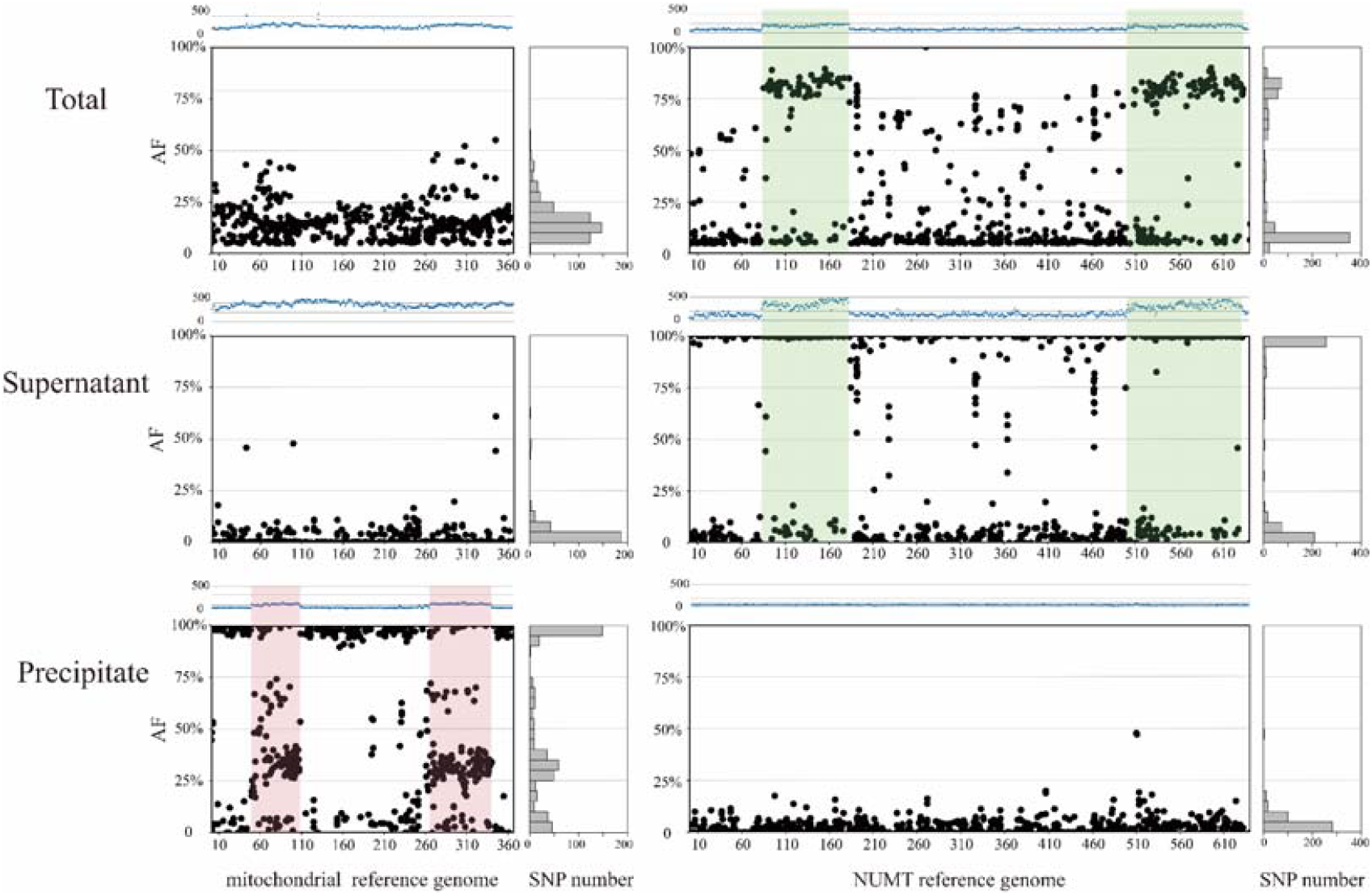
SNP calling and allele frequency (AF) calculation of NGS reads from total, supernatant, and precipitate DNA samples. From top to bottom, reads of total DNA, supernatant and precipitate DNA samples were mapped to the mitochondrial genome (left) and the NUMT sequence (right). The black scatter diagrams show the allele positions and frequencies of the SNPs. The blue scatter diagrams at the top of each figure represent the read depth. The total number of SNPs for every 5 % of allele frequency is shown by the bar plots on the right side of each figure. Red and green highlights correspond to Fig. 2.

### The CpG methylation status of different tissues and through the time course of growth in Col-0 mitoDNA

A similar RE and amplification assay was carried out on DNA from different tissues of different developmental stages of growth in Col-0. No mitoSNP showed through time course under *Pvu*I treatment and no numtSNP showed through time course under McrBC digestion, indicating the absence of mitoDNA methylation at least on the *rps3* gene area across various *Arabidopsis* Col-0 tissues and growth stages tested here (Fig. 4). In terms of Col-0 leaf mitoDNA in the orange frame of Fig. 4, the results of sanger sequencing indicate that within the group without restriction enzymes, mitoSNPs were predominantly detected in the first 30 days. As the leaves age, both types of SNPs emerge. This implies that the mitochondrial copy number ratio in young leaves may exceed that in older leaves, resulting in a very low proportion of numtSNPs that cannot be detected in young leaves (Preuten *et al*., 2010, Zhang *et al*., 2023). Taken together, no CpG methylation change was detected through the time course of DNA extracted from different tissues in Col-0 in our conditions.

**Fig. 4.**
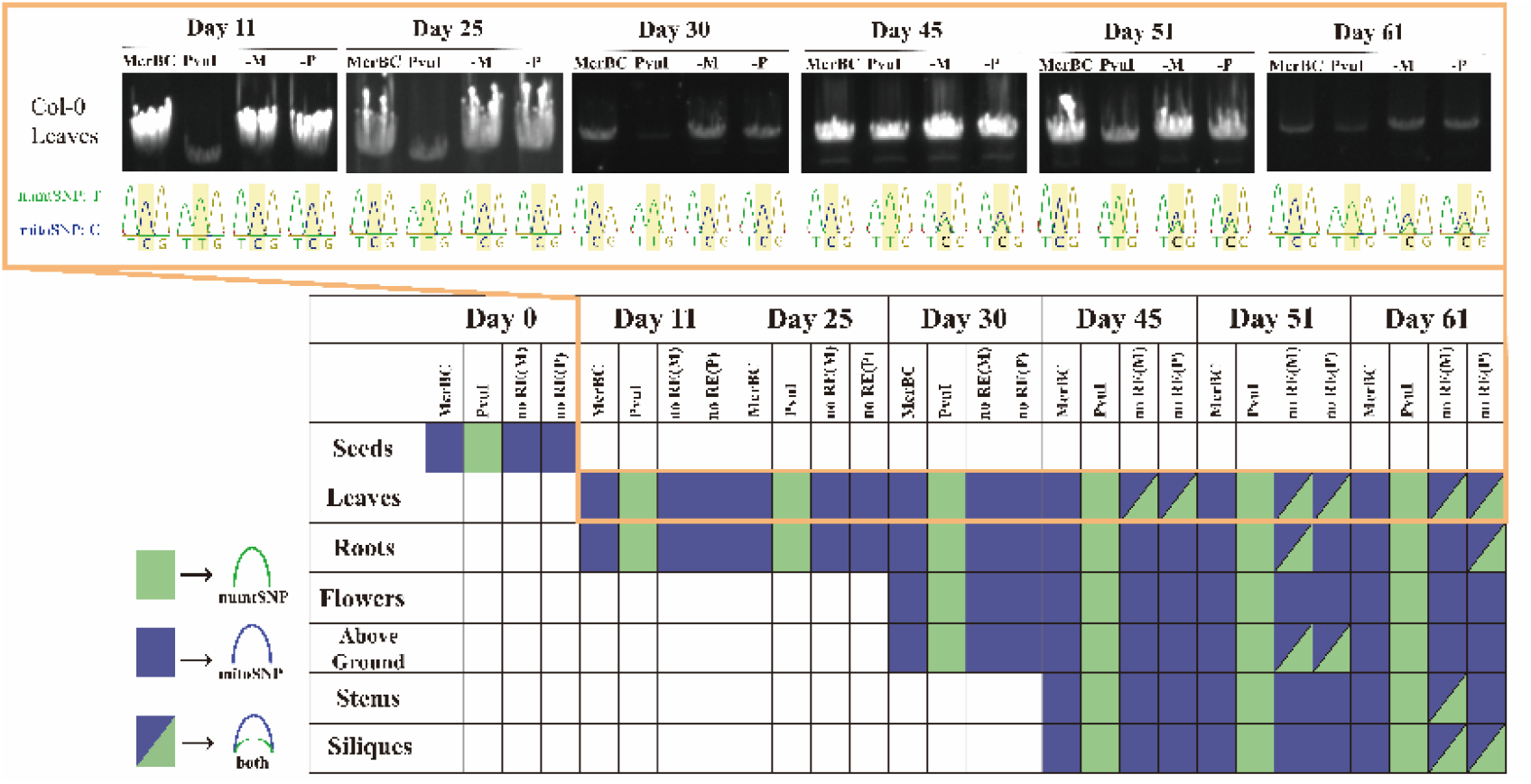
Sanger sequencing results of the *rps3* area amplification with DNA extracted from different tissues and through the time course of growth. For the picture in orange frame on the top, the base in the middle (yellow shade) represents the SNP sites. McrBC, CpG-methylation dependent restriction enzyme. *Pvu*I, CpG-methylation sensitive restriction enzyme. numtSNP: the type of SNP on NUMT; mitoSNP: the type of SNP on mitochondria. -M: No McrBC control, only the buffer of McrBC was used; -P: No *Pvu*I control, only the buffer of *Pvu*I was used. The green rectangle represents the numtSNP and the blue rectangle represents the mitoSNP. The white rectangles mean no data / not examined. The half-blue and half-green rectangles convey the existence of both SNPs.

## Discussion

Plant mitochondrial DNA methylation has been assessed before, mainly relying on the methylation-sensitive restriction endonucleases to detect the methylation of mitochondrial DNA of wheat embryos and carrot (Bonen *et al*., 1980, Simkova, 1998). Bisulfate sequencing has been considered the gold standard for detecting DNA methylation (Frommer *et al*., 1992, Gu *et al*., 2010, Muniandy et al., 2020), which involves the fragmentation of long DNA before sequencing. Additionally, the linearization and ultrasonic fragmentation prior to bisulfite conversion were shown to influence the final assessment of human mitochondrial DNA methylation levels (Guitton *et al*., 2022). Meanwhile, a study found that even in artificially synthesized unmethylated DNA, the conversion rate from cytosine to uracil after bisulfate treatment still cannot reach 100% (Matsuda et al., 2018).

Col-0 is still the golden standard ecotype of the model plant, *Arabidopsis thaliana*, widely used also in organelle studies. For Col-0, which possesses a NUMT larger than the entire mitochondrial genome with high identity, which introduces significant interference in determining DNA methylation in the mitochondrial genome. Fortunately, the high-quality nucleotide sequence of the NUMT on chromosome 2 serves as a significant rescue to carry out the methylation study because we could get the information of SNPs between the mitochondrial and the almost identical NUMT sequences (Fields *et al*., 2022b). In this study, the restriction enzyme check suggested a lack of CpG methylation in Col-0 mitochondrial DNAs, and the high CpG methylation of corresponding nuclear genome NUMT of *rps3* area. The rudimentary appraisal through two kinds of restriction enzymes on DNA from different tissues and different developmental stages of Col-0 also failed to detect CpG methylation on the *rps3* gene in the mitochondrial genome. In plants, cytosine methylation usually methylated at CG, CHG, and CHH context (Lister et al., 2008), yet the utilization of MBD protein in this study limited the examination of methylation within CG context. NGS sequencing further validated that this affinity precipitation method can effectively separate mitochondrial DNA from NUMTs to a significant extent via the difference in cytosine methylated level in the whole length. It was observed that MBD protein preferentially binds to CpG methylated areas with high-density (Zhang *et al*., 2006). The result of this study indicated that the MBD protein does not bind to DNA from mitochondria. Therefore, it is suggested that there may be no high-density CpG methylation in the mitochondrial DNA of Col-0. Moreover, this result also conveys the high CpG methylation in NUMTs, as reported before (Fields et al., 2022b). As a gene editing tool (Bogdanove and Voytas, 2011, Moscou and Bogdanove, 2009), transcription activator-like effectors (TALEs) are known for their sensitivity to cytosine methylated sequences (Bultmann *et al*., 2012, Kaya *et al*., 2017). Recent findings with mitoTALENs indicate their high efficiency in editing the mitochondrial genome (Arimura *et al*., 2020, Kazama *et al*., 2019, Forner *et al*., 2023), combined with this study, indicating this heightened editability may attributed to the low methylation status of mitochondrial DNA. Taking all the results into account, we suggest that there is absence of CpG methylation on Col-0 mitochondrial DNA.

## Materials and Methods

### Plant growth conditions and DNA extractions

The *A. thaliana* Col-0 was disinfected for 20 min with a solution (9% sodium hypochlorite, 2% Triton) and washed with sterile water. Then, evenly dispense onto 1/2 MS medium with 1% agarose and vernalize at 4°C in the dark for 7 days. After vernalization, the plants were transferred to the incubator for 14 days, then transplanted in Jiffy pots (Jiffy-7® Peat Pellets, http://www.jiffypot.com/) and cultivated under 22°C, 50-150 μmol m^−2^sec^−1^ light and 16 hours light/8 hours dark conditions until a size suitable for DNA extraction on 30 days after vernalization. DNA was extracted through the DNeasy Plant Pro kit and Maxwell RSC Plant DNA kit following the manufacturer’s instructions.

### Restriction enzyme treatment

Methylation-dependent restriction enzyme McrBC from TaKaRa (Code:1234A) and methylation-sensitive restriction enzyme *Pvu*I from NEW ENGLAND BioLabs (R3150S) was used, with reaction system following the manufacturer’s instructions. DNA was treated under 37°C for 90 min. After that, temperature was raised to 80°C for 20 min to inactivate enzymes.

### PCR analyses

PCR was carried out with KOD One® PCR Master Mix -Blue- (TOYOBO), amplified on *rps3* gene area, using the primers: Fw 5’-CACTGAGGGGAAGGTTGGTC-3’ and Rv 5’-GCTTTGCTACCGGGCTTCTA-3’. DNA after restriction enzyme treatment was used as PCR templates. PCR program was set according to the primer Tm and product size.

### MBD protein-based affinity precipitation

The MBD protein-involved immunoprecipitation was applied through NEBNext Microbiome DNA Enrichment Kit by following the manufacturer’s instructions.

### Next-generation sequencing and data analysis

Illumina’s next-generation sequencing platform was applied for the whole genome sequencing of total DNA, precipitated, and supernatant DNA samples. Low-quality and adapter sequences were trimmed from the raw reads using the fastp (Chen *et al*., 2018). We mapped the trimmed reads to the mitochondrial genome and the NUMT region, respectively, using the bwa-mem2 (https://github.com/bwa-mem2/bwa-mem2), and filtered out inadequately mapped reads with mapping identities ≤97% or alignment coverage rates ≤80% using samtools (https://github.com/samtools/samtools). SNP calling was completed by vcftools (https://github.com/vcftools/vcftools). Data visualization was performed using the ‘ggplot2’ package from R. Figure layout was conducted using Adobe Illustrator. Sequence alignments between the mitochondrial genome and the NUMT region were obtained and visualized using mummer4.

## Acknowledgements

We thank the technical assistance of Reiko Masuda for taking care of the plants and we are grateful to all the suggestions provided by our group members.

## Author contributions

Y.Y.Z. conducted the experiment and wrote the manuscript draft; M.O. and Y.Y.Z. analyzed and visualized the NGS data. S.A. and N.T. obtained the funding; S.A. conceived and supervised the whole study, finalizing the manuscript draft with the contributions from all authors.

